# Benchmarking DNA Foundation Models: Biological Blind Spots in Evo2 Variant-Effect Prediction

**DOI:** 10.64898/2026.03.10.710786

**Authors:** Vihaan Mathur, Ravi Sachidanandam

## Abstract

DNA foundation models such as Evo2 and DNABERT-2 show promise for variant-effect prediction (VEP), yet rigorous benchmarks assessing their ability to capture established biological constraints remain limited. Here, we introduce controlled benchmarks based on well-characterized nuclear and mitochondrial sequence features. Although Evo2 achieves strong aggregate performance on mitochondrial variants (AUROC = 0.927), our analyses reveal systematic blind spots in short-range signals, including codon usage bias, as well as spurious sensitivity to biologically neutral sequence context. Notably, changes in flanking sequences can cause pathogenic tRNA variants to be misclassified as benign. These findings suggest that Evo2’s predictive performance may rely more heavily on learned sequence regularities than on the biological constraints governing variant function, raising concerns about the clinical utility of zero-shot genomic foundation models. Our framework provides a principled basis for standardized model evaluation and for guiding the development of architectures better aligned with biological constraints.

## 1 Introduction

As the first draft of the human genome neared completion[1, 2], attention increasingly turned to understanding how differences between individual genomes might explain phenotypic variation[3]. At the same time, annotation of genomic elements, their functions and roles across tissues and cell types, became a central focus[4].

Over the past two decades, both variation data and functional annotations have accumulated in numerous public databases, supported by a wide array of software tools for exploration and interpretation. These resources represent the combined experimental and analytical efforts of many research groups worldwide.

Despite the scale of these efforts and the development of diverse computational approaches, including homology-based annotation[5], k-mer–based taxonomic methods[6], and machine-learning frameworks[7], we remain limited in our ability to reliably predict the functional consequences of genomic variation. This persistent gap has fueled interest in large DNA foundation models such as DNABERT-2[8] and the Evo series[9], which promise to extract biologically meaningful signals directly from raw sequence context that may not be readily apparent to traditional approaches.

Large genomic language models are trained on vast collections of DNA sequences from diverse species and learn statistical regularities from sequence context, functioning analogously to large natural language models, but operating on nucleotides rather than words.

Evo2[9] is a recently developed foundational DNA model trained autoregressively on the OpenGenome2 dataset (approximately 16, 000 eukaryotic genomes and totaling roughly 9 trillion bases), encompassing species across all domains of life. Given a raw DNA sequence as context, Evo2 can either predict the subsequent nucleotides or assign probability scores to a given sequence. Evo2’s hybrid convolutional framework, StripedHyena 2[10], departs from the standard transformer-based architectures[11] that dominate the language model space currently. It integrates short-, medium-, and long-range contextual “lenses,” enabling it in principle to capture biological features across multiple spatial scales—from codon-level motifs to exon–intron structure to chromosome-scale organization.

The model claims utility across a broad range of generative and evaluative tasks, including chromosome and genome design as well as variant-effect prediction. In particular, we focus on the three broad capabilities

Evo2 claims to achieve: 1) Generation of genome-scale DNA sequences, 2) Inherent capture of biological features across species, and 3) Zero-shot variant pathogenicity prediction (without explicit training on labelled variants) that surpasses state-of-the-art models.

The third claim is supported by their evaluation of a BRCA1 SNV dataset, where the 7B base model achieved an AUROC of 0.891[9].

We test these claims by asking if Evo2’s sequence likelihoods capture biologically relevant genomic signals, including codon bias[12, 13], mutational impact[14] (benign versus deleterious), and annotations of pseudogenes[15, 16] and NUMTs[17]. Pseudogenes are non-functional gene copies, often processed (intron-free) relics of retrotransposition, that are selectively unconstrained and accumulate mutations over time[18]. NUMTs are the pseudogene equivalents of mitochondrial DNA (mtDNA).

We focus on well-annotated regions of both the nuclear genome and mtDNA to provide controlled and interpretable benchmarks. The mitochondrial genome is a particularly tractable test case: it is compact, intron-free, and exhaustively characterized, with pathogenic variants linked to well-defined clinical syndromes including Leber’s Hereditary Optic Neuropathy (LHON; e.g. m.11778G*>*A in MT-ND4), MELAS (m.3243A*>*G in MT-TL1), MERRF (m.8344A*>*G in MT-TK), and Leigh syndrome (e.g. m.8993T*>*G in MT-ATP6). This rich annotation allows us to construct variant sets with known ground truth and to test model predictions against established biological expectations.

Variant-effect datasets are frequently imbalanced, often containing substantially more benign than deleterious mutations, or far more non-coding than coding variants. In such settings, raw accuracy can give an overly optimistic view of performance and aggregate metrics such as AUROC can obscure practical limitations. Use of confusion matrices [19] (section S2) for subsets (of variants) can help identify shortcomings as well as strengths of different tools, as we show here.

We identify several biological signals that appear to escape Evo2’s awareness which lead to benchmarking metrics that may guide improvements in future foundation models for genomic interpretation. These results also identify weaknesses that need to be addressed before foundational models can be deployed in the clinic.

## 2 Evaluation Framework

A DNA foundation model claiming biological utility must have internalized the statistical and structural grammar of genomes. Genomic signals operate across a wide range of length scales, and a model’s blind spots may depend on scale. We therefore organize our benchmarks into short(single nucleotide), medium(10s of nucleotides), and long-range signals (long regions of the genome ranging from 100’s to millions of bases) (Figure 1).

**Figure 1.**
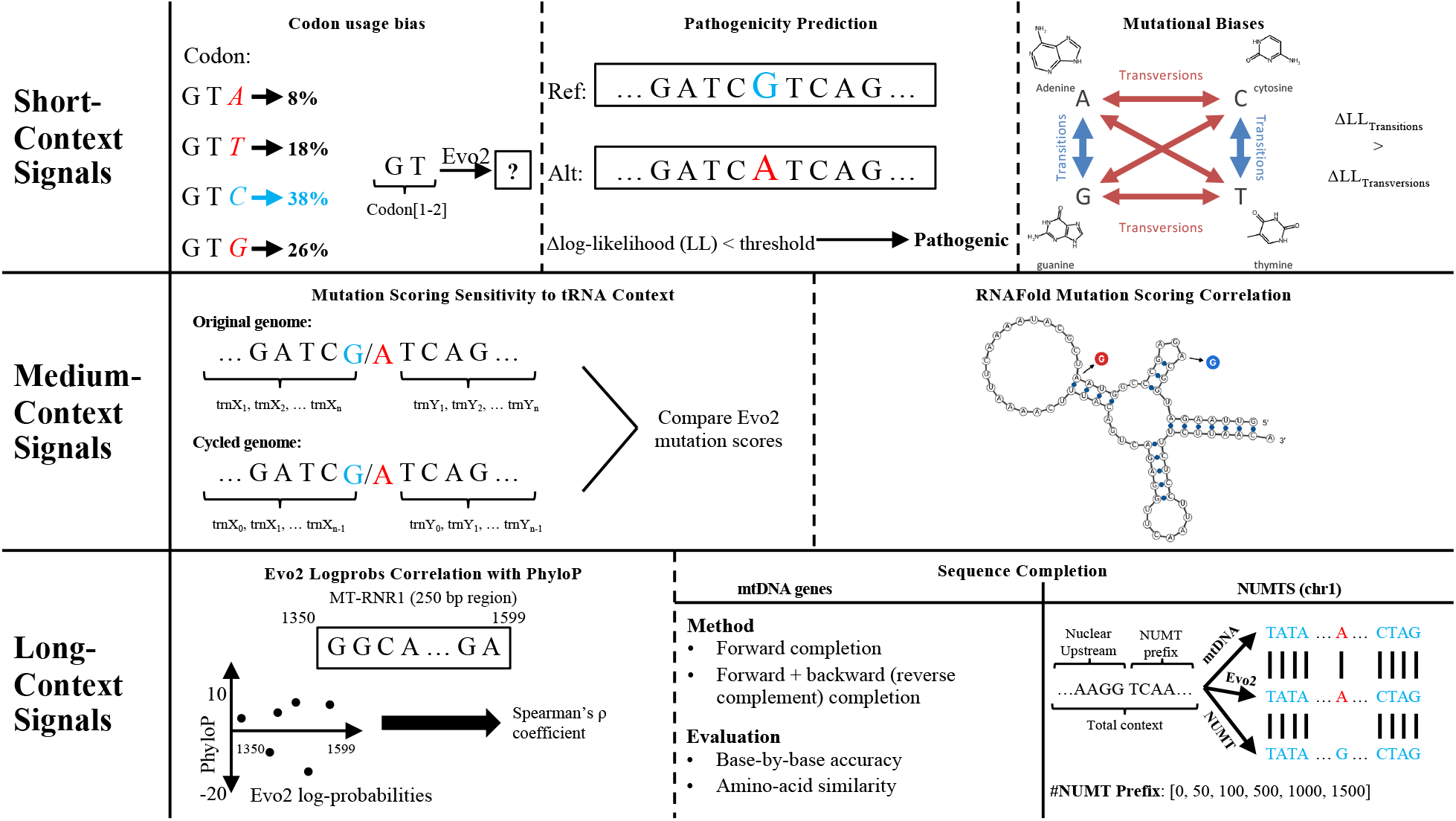
Schematic of framework for identifying genome-context awareness of foundational models at different length scales. Short-context signals include differential usage of synonymous codons (codon bias) and variant statistics, e.g. the relative abundance of transitions and transversions. Medium-context signals include variations that lead to destabilisation of tRNA structures and other structural features that involve interactions spanning 10’s to 100’s of nucleotides. Long-context signals can range from 1000’s to megabases including regions of conservation/homology.

### 2.1 Short-range Signals

#### 2.1.1 Single nucleotide variations

Changes at a single nucleotide can affect function through changes to protein sequence, regulatory elements, or chromatin architecture; those producing disease are pathogenic, others benign[20]. Foundation models provide sequence likelihood scores. Low scoring variants can be interpreted as being disfavored, indicating propensity for pathogenicity, but there isn’t an explicit identification of alleles as pathogenic. A key test of a foundation model is whether its sequence likelihood scores, computed without any variant-labeled training data, discriminate pathogenic from benign alleles. We evaluate this using a curated mitochondrial variant set spanning coding and non-coding regions.

#### 2.1.2 Transitions and Transversions

Nucleotides divide into purines (A, G; bicyclic) and pyrimidines (C, T; monocyclic). Transitions (A ↔ G; C ↔ T) are substitutions that preserve ring structure causing less steric distortion than ring-changing transversions (purine ↔ pyrimidine), and occur roughly three times more frequently in natural variation[14]. CpG sites add a further enrichment of C→T transitions through spontaneous deamination of methylated cytosine. A model that has internalized mutational biology should, at any given position, assign higher probability to the transition than to either transversion.

#### 2.1.3 Codon Usage Bias

The genetic code is degenerate: 61 sense codons specify 20 amino acids, so most amino acids are encoded by two to six synonymous codons, which usually differ at the third (wobble) position. Because wobble pairing lets one tRNA read several synonymous codons, cells carry fewer tRNA species than codons. Synonymous codons are not used equally[21, 22, 13], and two forces shape this bias:

- **Compositional bias**. Codon choice tracks the base composition of the surrounding genomic region. In GC-rich regions, G- and C-ending codons dominate; valine, for example, is encoded mostly by GUG and GUC rather than GUU or GUA. The GC-rich human gene *HSPA1A* shows this in extreme form. In mammals, this is the main source of gene-to-gene variation in codon usage.
- **Translational selection**. In fast-growing organisms, highly expressed genes favor codons read by abundant tRNAs[12]. In yeast, highly expressed genes encode leucine with UUG about 80% of the time and with CUA less than 1% of the time[23]. These preferences are species-specific: human and *E. coli* favor different arginine codons; of the six possible codons for arginine (CGU, CGC, CGA, CGG, AGA, and AGG) humans heavily favor the AGA and AGG triplets (40% of the arginine in humans but only 4% in E.coli), while E. coli strongly prefers CGU and CGC (80% of arginine) which is why human genes are often recoded for bacterial expression[24, 25].

Codon bias therefore operates at two levels: a genome-wide preference for each species, and a gene-level preference set by local composition. A model that has learned coding sequence should reproduce both. Given the start of a strongly biased gene, it should continue with that gene’s preferences rather than the genome average. We test this by having Evo2 complete coding sequences and comparing its synonymous codon choices with uniform, genome-wide, and gene-local usage.

#### 2.1.4 Mitochondrial Codon Idiosyncrasies

Vertebrate mtDNA uses a post-endosymbiotic variant genetic code with three critical distinctions: UGA encodes tryptophan, AUA encodes methionine, and AGA/AGG serve as stop codons (terminating human *MT-CO1* and *MT-ND6*).

Start and stop signals are notably flexible compared to nuclear genes. Human mitochondrial translation initiates with AUG (9 genes), AUA (3), or AUU (*MT-ND2*), all decoded as initiator methionine (Supplementary Table S4), while other vertebrates utilize additional initiators such as AUC[26, 27]. For termination, six genes rely on post-transcriptional polyadenylation to complete truncated U or UA codons into UAA[28, 29]. Clinical and population data confirm this permissiveness: mutations that disrupt canonical initiation are often benign, including substitutions in *MT-ND1* (at 3307/3308 which affect AUA)[30] and the haplogroup F2-defining m.12338T*>*C (AUA → ACA) in *MT-ND5* [31].

Consequently, substitutions at valid mitochondrial start codons (e.g., AUG↔AUA) or stop codons (e.g., UAA→UAG) should be well-tolerated. A model defaulting to nuclear rules, however, would flag them as deleterious or misinterpret terminating AGA/AGG codons as missense arginine. Because mitochondrial sequences are abundant in public databases, a foundation model trained at Evo2’s scale should in principle learn this variant code; we test whether Evo2 recognizes these rules or defaults to nuclear conventions.

### 2.2 Medium-range Signals

#### 2.2.1 tRNA Sensitivity to Flanking Context

Transfer RNAs fold into a conserved cloverleaf structure[32] whose aminoacylation and decoding properties—and the pathogenic mutations disrupting them, such as m.3243A*>*G in *MT-TL1* and m.8344A*>*G in *MT-TK* —are governed strictly by the tRNA’s internal sequence rather than its genomic neighborhood (Supplementary S1), a locality captured by structure-based tools like RNAfold[33]. Consistent with this independence, mitochondrial tRNAs undergo extensive, selectively neutral positional rearrangement across species[34]. To test whether Evo2 inappropriately incorporates flanking sequence into tRNA variant scores, we cyclically permuted mitochondrial tRNA genomic positions while keeping internal sequences intact; substantial shifts in predicted pathogenicity under this positional rearrangement would reveal spurious context sensitivity.

### 2.3 Long-range Signals

#### 2.3.1 Gene Completion

A generative genomic model should reconstruct missing gene segments by leveraging learned coding structure alongside flanking sequence. Because mitochondrial genes are compact and intron-free, masked completion of internal gene segments directly tests whether a model captures long-range coding dependencies rather than relying on local sequence statistics.

#### 2.3.2 NUMTs and Contextual Disambiguation

Nuclear mitochondrial DNA segments (NUMTs) are pseudogenic mtDNA fragments integrated into the nuclear genome[17]. Under relaxed selective constraint, NUMTs accumulate mutations that diverge from the extant mitochondrial reference, providing a natural test for genomic disambiguation. A biologically grounded model provided with sufficient nuclear context should distinguish NUMTs from authentic mtDNA and treat NUMT-specific variants as benign. We evaluate whether Evo2 recognizes these genomic compartments; defaulting to the mitochondrial sequence despite nuclear context would indicate that its variant effect scores are confounded by sequence similarity to mtDNA.

#### 2.3.3 Correlation with Evolutionary Conservation

Evolutionary conservation reflects purifying selection and functional constraint. A model whose likelihoods capture biological fitness should assign higher log-probabilities to conserved positions. We test this by correlating Evo2 per-base log-probabilities with PhyloP scores[35, 36] across the mitochondrial genome.

## 3 Methods

### 3.1 Mitochondrial Variant Dataset

We constructed a labeled mitochondrial variant dataset consisting of pathogenic and benign mutations. Variant counts and sources for every analysis in this study are summarized in Table S5.

**Pathogenic variants** (n = 135) were obtained from MITOMAP[37] and restricted to confirmed disease-causing mutations, and from ClinVar[38], restricted to entries labeled pathogenic or likely pathogenic.

**Benign variants** (n = 652) were obtained from the ClinGen frequency dataset and ClinVar [38], restricted to entries labeled benign with high-confidence review status (either reviewed by expert panel, or is a practice guideline, or a criteria provided by multiple submitters with no conflicts). All variant types were included (substitutions, insertions, deletions, and multi-base indels; Table S5).

The mitochondrial reference is the revised Cambridge Reference Sequence (NC 012920.1); nuclear coordinates use GRCh38.p13/hg38.

### 3.2 Evo2 Inference and Scoring

#### 3.2.1 Sequence generation

Sequence completion (gene completion and NUMT continuation) was performed by querying NVIDIA’s hosted Evo2 40B checkpoint (`evo2-40b`) via the NVCF `/generate` endpoint with greedy decoding (`top k = 1`, `top p = 1.0`, `temperature = 1.0`) to ensure determinism. When required, logits or sampled probabilities were requested to obtain internal confidence scores.

#### 3.2.2 Log-likelihood computation

Given sequence *W* of length *L*, Evo2 logits

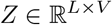

(*V* = 512 byte-level vocabulary) were obtained via the `/forward` endpoint with `“unembed”` output. Let (*x*_0_, …, *x*_*L−*1_) denote the byte-tokenized input.

Next-token log-probabilities were computed as:

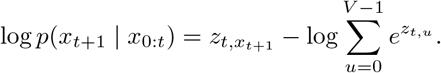

Mean log-likelihood (MLL) was defined as:

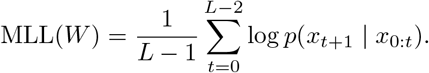

### 3.3 Codon Usage Bias Evaluation

The Effective Number of Codons (*N*_*c*_ or ENC) is a summary statistic that measures how much the codon usage of a gene departs from the equal use of synonymous codons[39]. ENC values range from 20 (extreme bias) to 61 (unbiased, uniform usage of codons).

#### Gene selection and generation

We evaluated three single-exon human genes (300–1,000 codons) spanning a spectrum of ENC (Table 1): *HSPA1A* (extreme bias, 642 codons), *ACTBL2* (moderate bias, 377 codons; chosen for lower GC_3_, the G/C bias in the third nucleotide of the codon, to decouple bias strength from GC content), and *OR8U1* (low bias, 310 codons). Conditioned on 128 upstream bases and the initial 40 codons, Evo2 generated the remaining in-frame coding sequence up to 333 codons (temperature = 1.0, top k = 4, top p = 1.0; 20 independent seeds pooled). To evaluate translational selection across taxa, the same protocol was applied to two highly expressed, strongly biased genes each (selected by Codon Adaptation Index[22]) in *S. cerevisiae* (*PDC1*, ENC 24.7; *PGK1*, ENC 26.6; SGD S288C R64-5-1) and *E. coli* (*tufA*, ENC 28.8; *groL*, ENC 28.4; K-12 MG1655, RefSeq NC 000913.3), generating their complete remaining sequence.

**Table 1.** Benchmark genes evaluated for codon usage bias generation.

| **Species** | **Gene** | **Codons** | **ENC** | **Selection basis** |
|---|---|---|---|---|
| *H. sapiens* | *HSPA1A* | 642 | 30.2 | Extreme bias (ENC) |
|  | *ACTBL2* | 377 | 47.3 | Moderate bias (ENC; low GC<sub>3</sub>) |
|  | *OR8U1* | 310 | 60.2 | Low bias (ENC) |
| *S. cerevisiae* | *PDC1* | 563 | 24.7 | High expression (CAI) |
|  | *PGK1* | 417 | 26.6 | High expression (CAI) |
| *E. coli* | *tufA* | 394 | 28.8 | High expression (CAI) |
|  | *groL* | 548 | 28.4 | High expression (CAI) |

#### Statistical comparison

For each amino acid, generated synonymous codon counts (*O*) were compared against three references (*f*): a uniform distribution, the gene’s observed full coding sequence (gene-local), and genome-wide usage[13]. We evaluated goodness-of-fit via *χ*^2^ tests against uniform and gene-local expectations rescaled to the observed count,

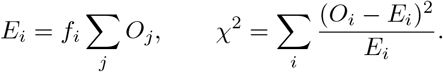

Because the *χ*^2^ statistic diverges when *E*_*i*_ = 0, amino acids containing codons absent from the endogenous sequence (*f*_*i*_ = 0) were excluded from the gene-local test.

To quantify divergence from gene-local and genome-wide frequencies without zero-count sensitivity, we computed the total-variation (TV) distance across the synonymous codons of each amino acid,

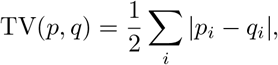

representing the minimum probability mass required to match the reference distribution. Final per-gene TV metrics were computed as the count-weighted mean across all 18 degenerate amino acids.

### 3.4 Mitochondrial Pathogenicity Prediction

For each variant at position *p* with reference allele *r* and alternate allele *a*, context windows were constructed:

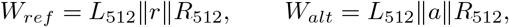

where 512 bases were included upstream and downstream. Mean log-likelihoods were computed:

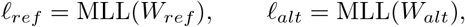

and the variant disruption score defined as:

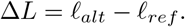

More negative Δ*L* indicates stronger disruption. A variant was called pathogenic when Δ*L* ≤ *τ* for threshold *τ*.

The dataset was split into calibration (70%) and test (30%) sets stratified by variant classification. Threshold selection used the Youden index[40]:

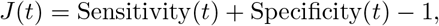

computed from the ROC curve. The threshold maximizing *J* on the calibration set was applied to the test set.

Performance metrics included sensitivity, specificity, AUROC, balanced accuracy, and confusion matrices (section S2).

#### 3.4.1 Context-length sensitivity

We repeated variant effect prediction on the same set at 64, 128, 256, 512, 1024, 4096, and 8192 bp flanking contexts without a species tag. A new Youden threshold is calibrated on the training set of each flank and applied to the test set (Table 5).

### 3.5 Mitochondrial Start/Stop Codon Interchangeability

For each mitochondrial protein-coding gene, valid alternative mtDNA start and stop codons were considered only when they differed from the annotated codon by a single nucleotide. Each retained substitution was scored using the same Δ*L* procedure as the pathogenicity analysis (128bp per flank) and classified using the Youden-optimal threshold derived from the calibration set at the same flank size (Table 5).

### 3.6 Transitions vs. Transversions

We ran Evo2 once over D-loop positions 1-500, with 512 bp of upstream context from the circular genome, and recorded its predicted probability for each base at every position. For each position, we checked whether the transition was the most probable of the three non-reference bases and what probability share it had among the three non-reference bases.

### 3.7 Sensitivity to tRNA Context

Annotated mitochondrial tRNAs were cyclically permuted while leaving other genes unchanged. Variant scoring was repeated (1024bp per flank) and confusion matrices compared between original and permuted genomes. To determine if changes in variant classification were statistically significant, a two-sided McNemar test on the paired correct/incorrect outcomes was used.

### 3.8 Gene Completion Experiments

Mitochondrial genes from ten species (human, mouse, rat, chicken, frog, zebrafish, *Drosophila*, yeast, *Arabidopsis, C. elegans*) were evaluated. Every gene completion prompt was prefixed with a species-specific phylogenetic lineage tag generated from GBIF taxonomy (for human, |`D__ANIMALIA;P__CHORDATA;C__MAMMALIA;O__PRIMATES;F__HOMINIDAE;G__HOMO;S__HOMO SAPIENS`|). Three strategies were tested:

#### Forward completion

500 bp upstream plus the first two-thirds of the gene were provided; Evo2 generated the remaining third.

#### Masked middle with directional selection

Each gene was divided into thirds; the middle third was masked.

Forward: upstream context + first third. Backward: downstream context + last third (reverse complement).

Log-likelihood:

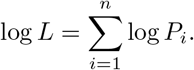

The higher-likelihood direction was selected.

#### Masked middle with base-level consensus

At each position, if forward and backward predictions agreed, the shared base was chosen. Otherwise, the base with higher log-probability exceeding a margin *m* was selected; ties were resolved using the direction with higher global log-likelihood.

Evaluation metrics included base-level accuracy and forward–backward win rates.

### 3.9 NUMT Generation and Divergence Analysis

A high-identity NUMT (GRCh38 chr1:629079–634924) was selected. Input sequences consisted of 1000 bp nuclear upstream context followed by NUMT prefixes, with prefix lengths of 0, 50, 100, 500, 1000, and 1500 bp. Evo2 generated 200 bp continuations following the prefix string.

Generated sequences were compared to the NUMT continuation and homologous mtDNA. At positions where NUMT and mtDNA diverged, Evo2 predictions were recorded to determine whether the model favored NUMT-specific or mitochondrial sequence identity.

### 3.10 Correlation with Evolutionary Conservation

A 250 bp region of the 12S rRNA gene (rCRS 1350–1599) was analyzed. Per-base Evo2 log-likelihoods were correlated with PhyloP conservation scores using Spearman’s *ρ* due to expected nonlinearity and non-normality.

## 4 Results

### 4.1 Codon Usage Bias

#### Compositional bias

Table 2 summarizes per-amino-acid *χ*^2^ tests on Evo2’s generated codon usage. The majority of amino acids differ from the null distribution, but also from the true gene’s distribution of synonymous codons. For example, the true leucine distribution in *HSPA1A* is 0%/8%/88%/0%/0%/4% (CTA/CTC/CTG/CTT/TTA/TTG), while Evo2’s generated distribution is 3%/12%/71%/5%/2%/7%. Evo2 produces CTA, CTT, and TTA, which are all absent from the true sequence, at nonzero frequency. For the two biased genes, generated usage sits closer to the gene’s own distribution than to genome-wide usage (HSPA1A: TV 0.113 vs. 0.307), while for the near-unbiased *OR8U1* the ordering reverses.

**Table 2.** Evo2’s generated codon usage compared against three reference distributions. ENC (Effective Number of Codons) ranges from 20 (maximal bias) to 61 (no bias). *χ*^2^ columns give the number of amino acids differing significantly (*p <* 0.05) out of those testable against each reference. Total-variation (TV) distance is the fraction of generated codons that would have to be reassigned to match the reference.

| **Gene** | **ENC** | χ<sup>2</sup> |  | Mean TV distance |  |
|---|---|---|---|---|---|
|  |  | vs. uniform | vs. gene-local | to genome-wide | to gene-local |
| *HSPA1A* (first 333 codons) | 30.2 | 18/18 | 7/8 | 0.307 | 0.113 |
| *ACTBL2* (first 333 codons) | 47.3 | 16/18 | 7/14 | 0.114 | 0.091 |
| *OR8U1* | 60.2 | 13/18 | 8/16 | 0.099 | 0.107 |

#### Translational selection

Table 3 compares true and Evo2-generated codon usage for two strongly biased genes. In *PGK1*, the true leucine codons are only TTG (88%) and TTA (12%). Evo2 reproduces TTG as the dominant codon but at a lower frequency (72%), and assigns 13% of leucine codons to the four CT-codons (CTA 8%, CTT 3%, CTG 1%, CTC 1%), none of which occur in the true gene. In *groL*, the true arginine codons are only CGT (77%) and CGC (23%). Evo2 again predicts CGT as dominant but of less frequency (65%), slightly overuses CGC (27%), and assigns 8% of arginine codons to CGA, AGA, AGG and CGG, which are absent from the true gene.

**Table 3.** Evo2’s generated codon usage in yeast and *E. coli*. Percentages within each amino acid for the true coding sequence and for Evo2’s generated continuation (20 seeds pooled; 745 generated leucine codons for *PGK1*, 385 generated arginine codons for *groL*).

| *S. cerevisiae PGK1*, leucine |  |  | *E. coli groL*, arginine |  |  |
|---|---|---|---|---|---|
| **Codon** | **True** | **Evo2** | **Codon** | **True** | **Evo2** |
| TTG | 88% | 72% | CGT | 77% | 65% |
| TTA | 12% | 15% | CGC | 23% | 27% |
| CTA | 0% | 8% | CGA | 0% | 3% |
| CTT | 0% | 3% | AGA | 0% | 2% |
| CTG | 0% | 1% | AGG | 0% | 2% |
| CTC | 0% | 1% | CGG | 0% | 1% |

### 4.2 Pathogenicity Prediction

#### 4.2.1 Overall performance

Applying the Youden-optimal threshold (Δ*L* = −0.0081) to the test set, Evo2 achieves a true positive rate of 90.2%, true negative rate of 84.6%, balanced accuracy of 87.4%, and AUROC of 0.927 (Figure 2). While pathogenic variants are enriched at more negative Δ*L* values, the distributions overlap substantially, reflecting the difficulty of zero-shot discrimination.

**Figure 2.**
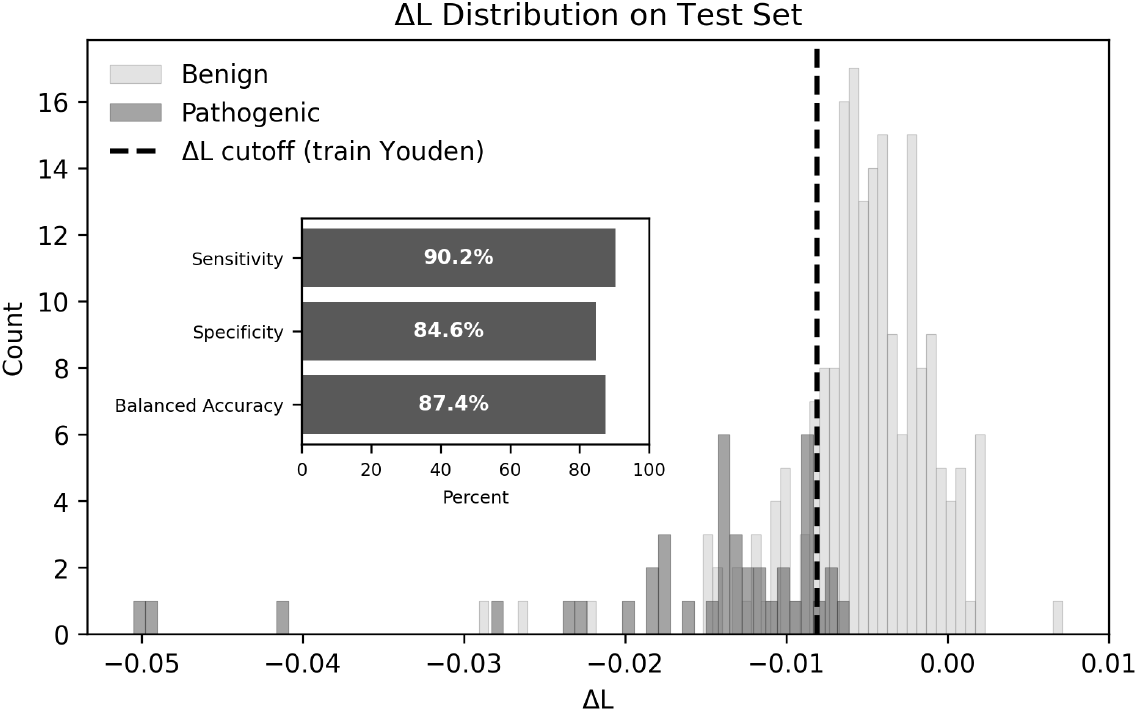
Evo2 zero-shot discrimination of mitochondrial variants. Distribution of Δ*L* = MLL(*W*_alt_) *−* MLL(*W*_ref_) for benign (blue) and pathogenic (orange) variants in the test set. More negative Δ*L* indicates greater disruption of sequence plausibility; pathogenic variants are expected to shift left. The dashed vertical line marks the Youden-optimal threshold (Δ*L* = *−*0.0081) derived from the calibration set. Pathogenic variants are enriched at more negative values, but with substantial overlap.

#### 4.2.2 Performance by region and variant class

Stratifying by genomic region and amino-acid consequence reveals important variability (Figure 3). For protein-coding missense variants, all 19 pathogenic variants were correctly identified, but 10 of 28 (35.7%) benign missense variants were misclassified as pathogenic. In RNA variants, sensitivity fell to 80% (a 20% false-negative rate), while 14.3% of benign RNA variants were incorrectly flagged. D-loop variants showed the poorest specificity, with 34.9% of benign variants called pathogenic.

**Figure 3.**
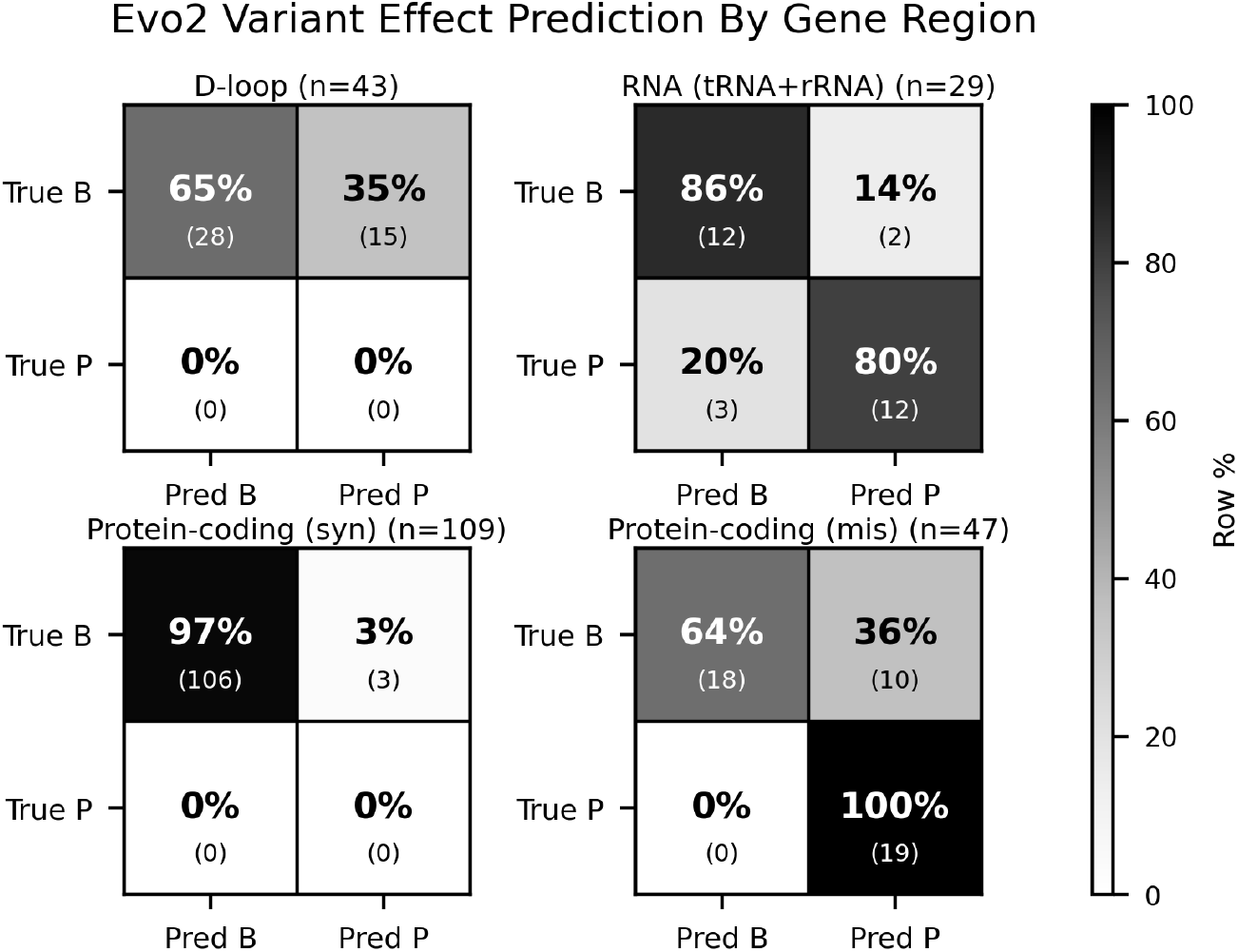
Evo2 performance stratified by genomic region and amino-acid consequence. Row-normalized confusion matrices for mitochondrial test-set variants, grouped by D-loop (non-coding), RNA genes (tRNAs and rRNAs), protein-coding synonymous, and protein-coding missense (228 of 236 test variants fall into these four categories; Table S5). Raw counts are shown in parentheses. Performance varies substantially across categories, with notable weaknesses in RNA variants and non-coding D-loop variants.

#### 4.2.3 Transitions vs. Transversions

Evo2’s predicted probabilities show a weaker preference for transitions than is seen in real variation (Table 4). Across the D-loop (1-500), the transition was the most probable non-reference base at 57.6% of positions, compared with 33.3% if all three substitutions were equally likely (*p* = 10^*−*28^, binomial test), and took 48.6% of the non-reference probability. In contrast, 93.2% of observed variants in this region are transitions. There was almost no preference at adenine positions (37.2%).

**Table 4.** Evo2’s preference for transitions across D-loop positions 1-500. For each reference base, the percentage of positions where the transition is Evo2’s most probable non-reference base, the mean share of non-reference probability assigned to the transition (33.3% for both if all substitutions were equally likely), and the percentage of observed variants in this region that are transitions.

| **Ref. base** | **Positions** | **Ts most probable** | **Ts prob. share** | **Observed Ts** |
|---|---|---|---|---|
| A | 145 | 37.2% | 33.6% | 92.9% (13/14) |
| C | 153 | 63.4% | 52.5% | 91.7% (11/12) |
| G | 69 | 72.5% | 57.3% | 87.5% (7/8) |
| T | 133 | 65.4% | 56.0% | 100% (10/10) |
| All | 500 | 57.6% | 48.6% | 93.2% (41/44) |

#### 4.2.4 Context-length sensitivity

Balanced accuracy was 0.81 at 64 bp and rose to a maximum of 0.87 at 512 bp. Beyond 512 bp, balanced accuracy and AUROC declined markedly, reaching 0.72 at 4096 bp and 0.73 at 8192 bp.

### 4.3 Mitochondrial start/stop codon interchangeability

For the three valid initiators (AUG, AUA, AUU), each initiator differs from the other two by a single base, giving two alternatives per gene across 13 genes; the seven genes with complete stops contribute one single-base alternative each. When presented with variants that swap one valid mitochondrial start codon for another (derived from the codons listed in Table S4), Evo2 classified all 26 alternatives as pathogenic (Table 6). For the set of alternative stop codons, Evo2 predicted 57.1% as pathogenic (4*/*7). Evo2 does not distinguish mtDNA start/stop codon usage from nuclear conventions.

**Table 5.** Pathogenicity performance by flanking context length. Balanced accuracy on the held-out test split (236 of the 787 variants) using a Youden threshold calibrated on the training split separately for each flank, and AUROC.

| **Flank (bp)** | **Balanced accuracy** | **AUROC** |
|---|---|---|
| 64 | 0.808 | 0.919 |
| 128 | 0.818 | 0.922 |
| 256 | 0.834 | 0.914 |
| 512 | 0.874 | 0.927 |
| 1024 | 0.801 | 0.883 |
| 4096 | 0.722 | 0.815 |
| 8192 | 0.727 | 0.797 |

**Table 6.** Evo2 predictions for mitochondrial start and stop codon-preserving variants. Despite maintaining valid mitochondrial start or stop codons based on the human mitochondrial genetic code, all the 26 start-codon preserving variants were called pathogenic by Evo2, while 4 out of 7 stop-codon preserving variants were predicted to be pathogenic by Evo2.

| **Variant Class** | **Total** | **Pred. Pathogenic** | **Pred. Benign** | **% Pred. Pathogenic** |
|---|---|---|---|---|
| Start codon preserving | 26 | 26 | 0 | 100% |
| Stop codon preserving | 7 | 4 | 3 | 57.1% |

Two factors temper this result. First, start and stop positions are strongly conserved, so a model may penalize any substitution there without reference to the genetic code; separating these requires comparing start-preserving against start-destroying substitutions. Second, the compact mitochondrial genome contains overlapping reading frames, and a subset of these variants alters the amino acid encoded in the overlapping gene (*MT-ATP6* /*MT-ATP8, MT-ND4* /*MT-ND4L*), where a pathogenic call may be correct.

### 4.4 tRNA Context Permutation

In the original genomic context, Evo2 correctly identified 48 of 72 pathogenic tRNA variants (sensitivity 66.7%), with a false-positive rate of 23.4% among benign variants. After cyclic permutation of all mitochondrial tRNA positions, leaving their sequences intact but changing their genomic neighborhoods, sensitivity collapsed to 19.4% (14/72 pathogenic variants identified), while specificity rose to 93.8% (Figure 4). Among pathogenic variants, 36 (out of 48, 75%) changed from correct to incorrect classification while only 2 changed in the opposite direction (McNemar’s test, *p* = 8.6 × 10^*−*8^).

**Figure 4.**
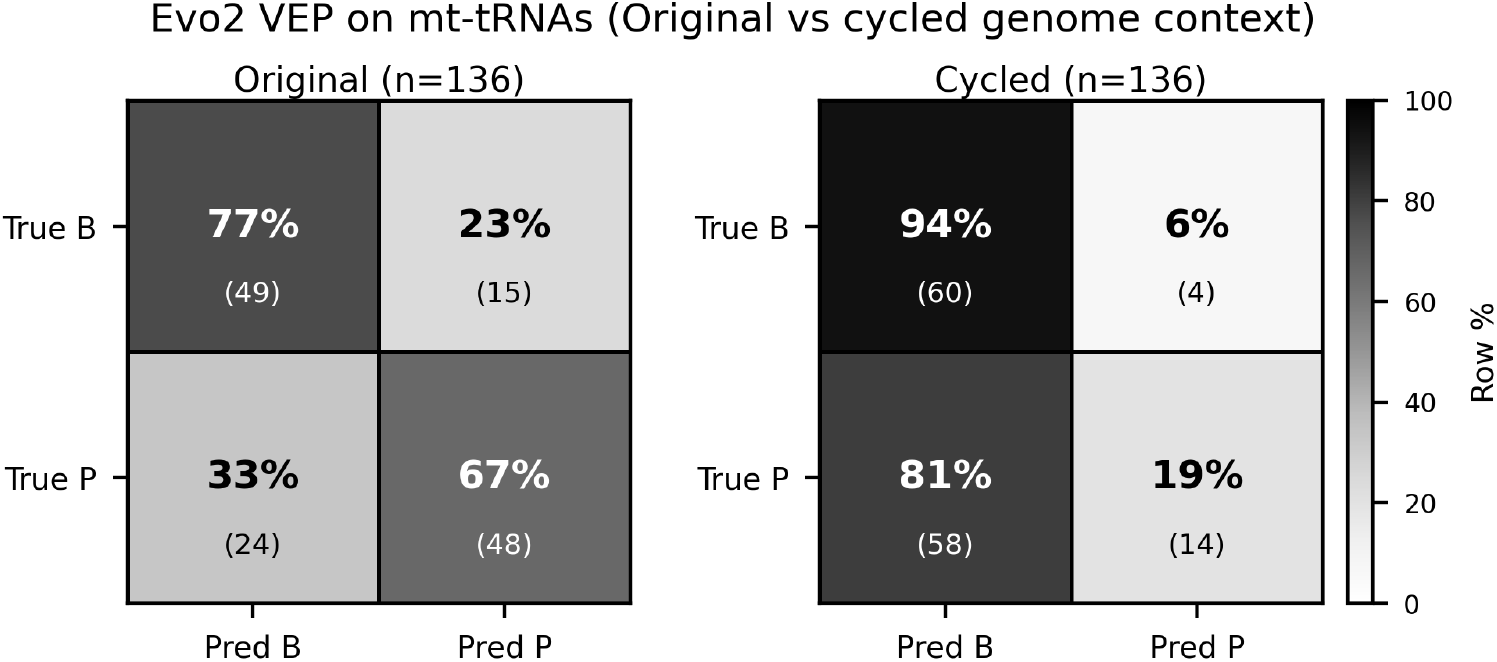
Effect of tRNA positional permutation on Evo2 pathogenicity predictions. Row-normalized confusion matrices for mitochondrial tRNA variants in the original genomic context (left) and after cyclic permutation of all 22 tRNA positions (right). tRNA internal sequences are identical in both conditions; only flanking context differs.

### 4.5 Gene Completion

Across all ten species and the three mitochondrial gene classes, Evo2 achieved a mean base-level completion accuracy of 86.2% (371 genes), highest for tRNAs (89.9%, mean over species), followed by mitochondrial OXPHOS genes (83.1%), and lowest for rRNAs (avg 65.2%) (Figure 5). Human mtDNA OXPHOS genes were completed more accurately (95.5%) than any other species (69.2% − 90.1%), including more closely related mammals such as mouse and rat, suggesting a training data bias rather than a reflection of conservation. Within humans, accuracy across OXPHOS complexes did not track known mutational constraint: Complexes I (ND1, ND2, ND3, ND4, ND4L, ND5, and ND6), IV (MT-CO1, MT-CO2, and MT-CO3), and V (MT-ATP6, MT-ATP8) all exceeded 95%, while Complex III(CYTB), which is severely disruption-intolerant[41], was the least accurately completed at 85.0%. Complex II has no genes in the mtDNA. Together these patterns suggest that Evo2’s completions are shaped primarily by sequence familiarity rather than biological constraint.

**Figure 5.**
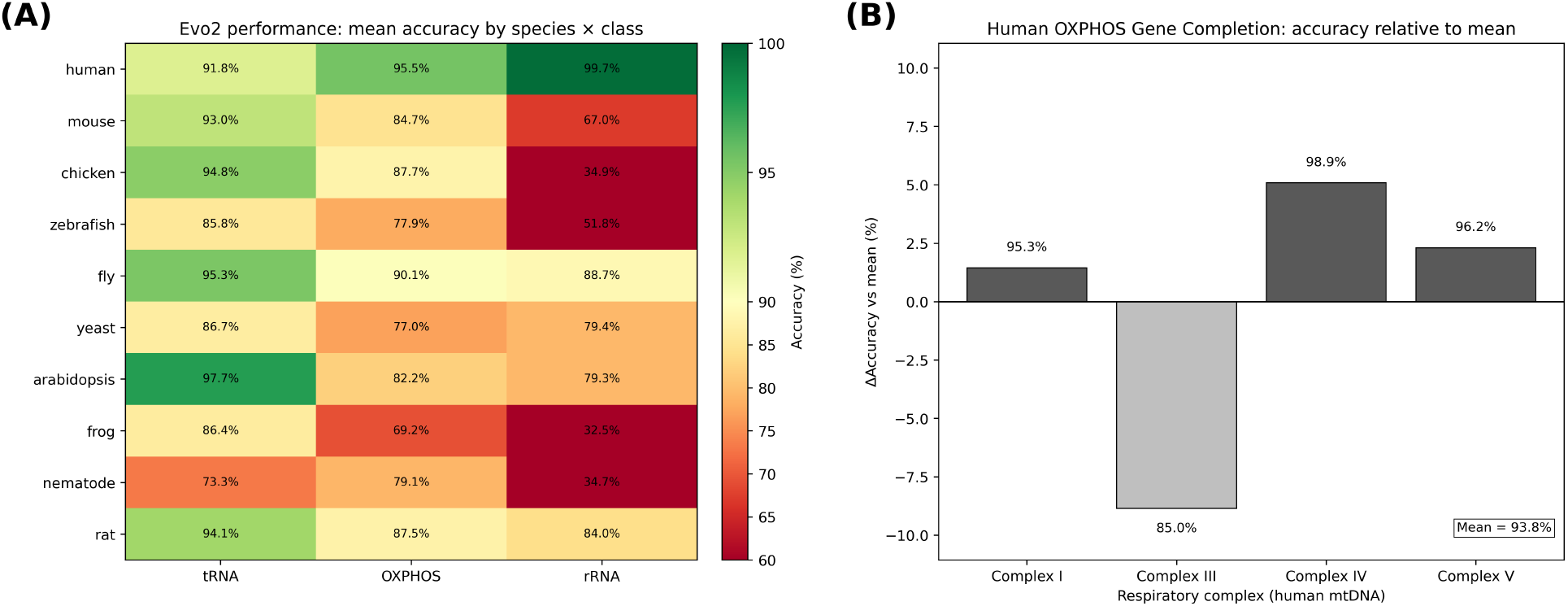
Evo2 base-by-base gene completion accuracy by gene class and species. (A) Mean completion accuracy for mitochondrial tRNAs, OXPHOS genes, and rRNAs across ten species. Human genes show the highest accuracy even when they are not the most evolutionarily conserved. Accuracy degrades in non-mammalian species (e.g. frog rRNAs: 32.1%). (B) Completion accuracy for human OXPHOS genes stratified by complex. The highly conserved Complex III (CYTB is the only mtDNA gene in this complex) shows the *lowest* accuracy (85.0%), while the remaining complexes all exceed 95% (Complex I 95.3%, Complex IV 98.9%, Complex V 96.2%; mean 93.8%).

For the masked-middle completion task, neither direction dominates overall (forward more accurate for 147 of 371 genes, backward for 139, tied for 85; Figure S2); the near-parity that is itself notable for an autoregressive model. The preferred direction instead varies by species, from backward-leaning (*Arabidopsis* 10/19, fly 6/13) to forward-leaning (human 18/9, nematode 21/13), indicating that downstream sequence carries information the forward pass does not recover. We therefore combine both directions in the consensus strategy.

### 4.6 NUMTs

To test whether Evo2 can distinguish authentic mitochondrial sequence from nuclear mitochondrial pseudogenes, we supplied variable-length prefixes of the high-identity NUMT (GRCh38, chr1:629,079-634,924) preceded by 1,000 bp of nuclear upstream context, and recorded Evo2’s 200 bp continuations.

Without any NUMT prefix (0 bp), Evo2 performs near chance (~28-29%) (Table 7). Accuracy rises sharply with prefix length, peaking at 500-1000 bp before degrading at 1500 bp. Notably, at every nonzero prefix length Evo2’s predictions match the mtDNA reference marginally better than the NUMT itself, indicating that the model treats the NUMT as if it were authentic mitochondrial sequence. This is confirmed by examining individual divergence sites: at positions where the NUMT has accumulated mutations relative to mtDNA, Evo2 predominantly predicts the mitochondrial allele (Tables 8, 9, 10).

**Table 7.** Evo2 continuation accuracy as a function of NUMT prefix length, compared against both the NUMT sequence and the homologous mtDNA.

| **NUMT prefix (bp)** | **Match with NUMT (%)** | **Match with mtDNA (%)** |
|---|---|---|
| 0 | 29.0 | 28.1 |
| 50 | 66.5 | 67.0 |
| 100 | 83.5 | 84.0 |
| 500 | 99.5 | 100.0 |
| 1000 | 97.0 | 99.5 |
| 1500 | 78.0 | 80.0 |

**Table 8.** Divergence sites between NUMT and mtDNA for prefix set at 500 bp. Evo2 preferentially picks the allele present in mtDNA over the correct one for genomic NUMT.

| **Index** | **chr1 position** | **chrM position** | **mtDNA** | **NUMT** | **Evo2** |
|---|---|---|---|---|---|
| 46 | 629,626 | 4,456 | C | T | C |

**Table 9.** Sites that differ between NUMT and mtDNA. Evo2, using a prefix of 1000 bp, matches the mtDNA allele at all five positions.

| **Index** | **chr1 position** | **chrM position** | **mtDNA** | **NUMT** | **Evo2** |
|---|---|---|---|---|---|
| 4 | 630,084 | 4,914 | C | T | C |
| 30 | 630,110 | 4,940 | C | T | C |
| 48 | 630,128 | 4,958 | A | G | A |
| 81 | 630,161 | 4,991 | G | A | G |
| 131 | 630,211 | 5,041 | T | C | T |

**Table 10.**
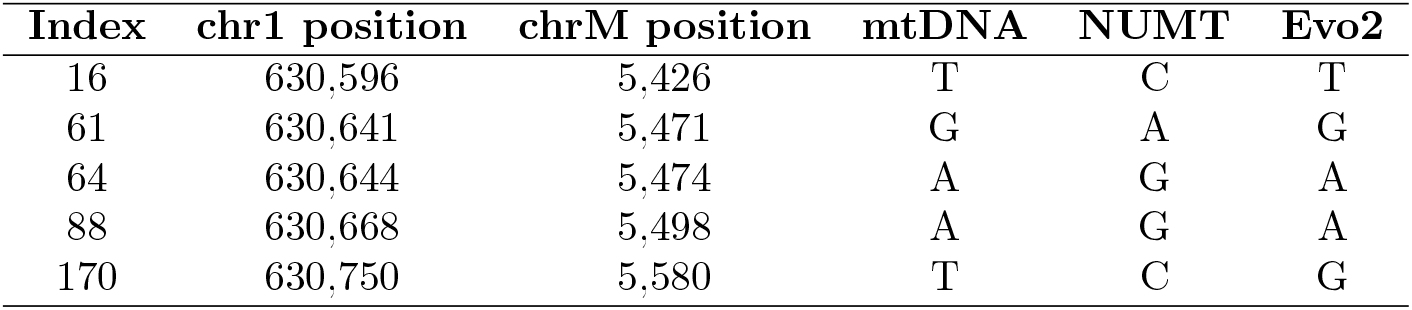
Sites differing between NUMT and mtDNA. Evo2, using a prefix of 1500 bp. matches the mtDNA allele in 4 of 5 cases.

| **Index** | **chr1 position** | **chrM position** | **mtDNA** | **NUMT** | **Evo2** |
|---|---|---|---|---|---|
| 16 | 630,596 | 5,426 | T | C | T |
| 61 | 630,641 | 5,471 | G | A | G |
| 64 | 630,644 | 5,474 | A | G | A |
| 88 | 630,668 | 5,498 | A | G | A |
| 170 | 630,750 | 5,580 | T | C | G |

### 4.7 Evolutionary Conservation

Across the 250 bp mt-RNR1 window, the Spearman correlation between Evo2 per-base log-probabilities and PhyloP conservation scores was *ρ* = 0.77, indicating a strong rank-level association. However, local conservation peaks do not correspond to peaks in Evo2’s log-probability, and sustained regions of conservation go largely undetected (Figure 6).

**Figure 6.**
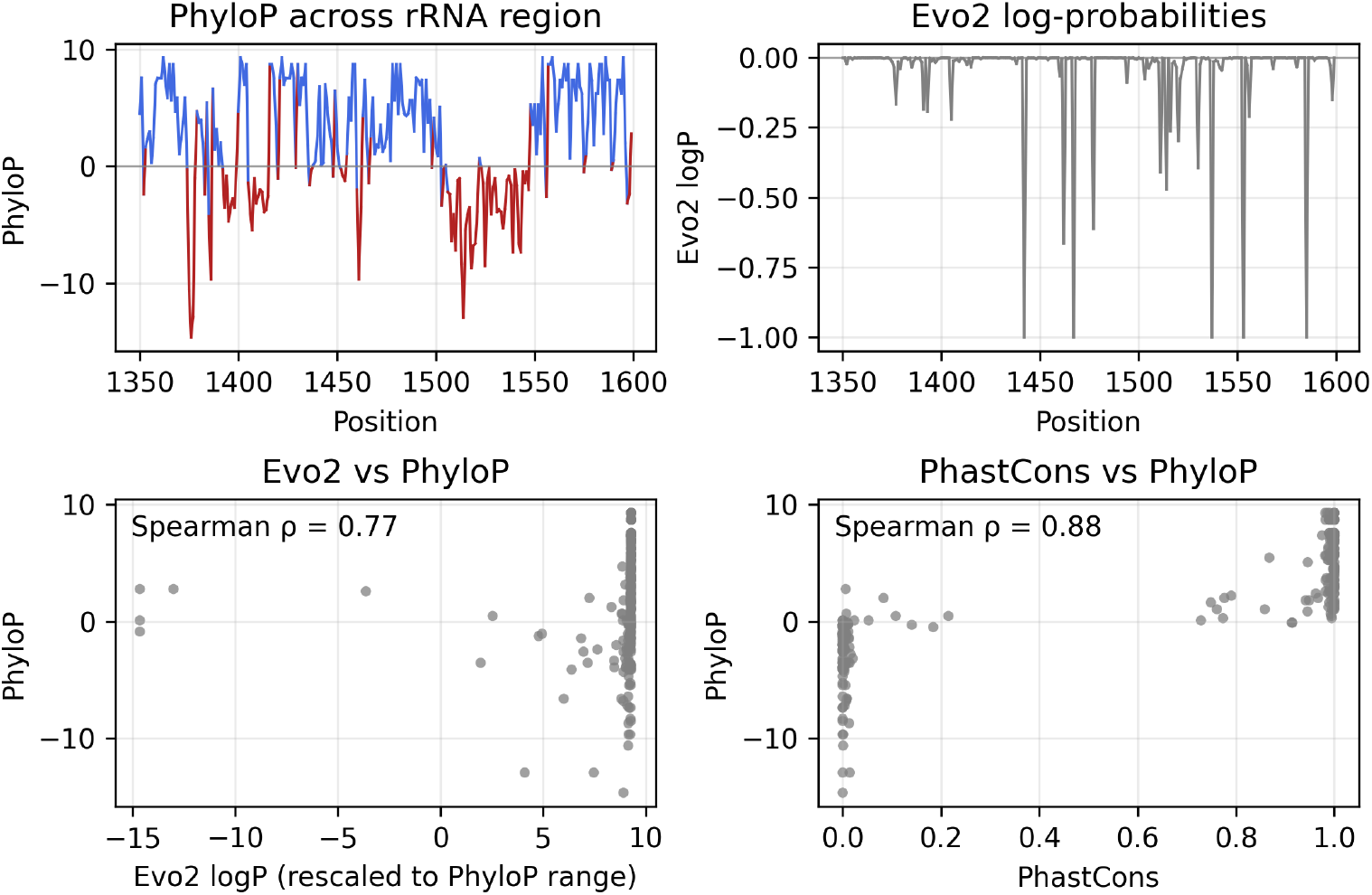
Evo2 log-probability vs. PhyloP conservation across the mt-RNR1 gene (rCRS 13501599). Top: PhyloP100way scores (positive = conserved, negative = evolving rapidly). Middle: Evo2 per-base log-probabilities. Bottom: Scatter plot of the two tracks. Evo2 has spikes that do not consistently correspond to local PhyloP peaks and sustained regions of conservation are not detected by Evo2.

## 5 Discussion

### 5.1 Performance Variability and Class Imbalance

Aggregate metrics mask substantial cross-category variation. Because benign variants predominate and certain regions (e.g., the non-coding D-loop) lack pathogenic entries entirely, global accuracy tracks the majority class. Consequently, strong aggregate AUROC obscures poor pathogenic detection (Supplementary S2), with false-positive rates exceeding 30% in several categories—a clinically consequential blind spot that mandates category-stratified evaluation across distinct mitochondrial functional classes.

Furthermore, Evo2’s transition preference is directionally consistent but quantitatively muted: across D-loop positions, it favors transitions at only 57.6% of sites compared to 93.2% in observed variants, and exhibits no transition preference at adenines.

### 5.2 Codon Usage Bias

Codon usage bias is a pervasive genomic signal that a model trained on trillions of eukaryotic nucleotides should readily capture. Yet Evo2 fails to reproduce the strong compositional bias against A/T-ending codons in *HSPA1A*, generating coding sequences where 11% of codons must be reassigned (total-variation distance ≈ 0.11) to match the endogenous distribution.

### 5.3 The Mitochondrial Genetic Code

The vertebrate mitochondrial code differs from the standard code at four codons and utilizes three distinct initiators. Despite ample representation in public genomic training data, Evo2 misclassifies every start-preserving substitution as pathogenic, though conservation at these sites partially confounds predictions (see Results).

Applying nuclear translation rules to mitochondrial sequence represents a fundamental compartment error analogous to mistaking NUMTs for authentic mtDNA. Because mitochondrial termini harbor substantial benign variation, this failure inflates false-positive rates precisely within clinically delicate regions.

### 5.4 Spurious Context Sensitivity: The tRNA Permutation Test

tRNA function—aminoacylation, modification, and decoding—is determined strictly by intramolecular interactions within the 70–100 nt molecule, independent of flanking sequence (Supplementary S1). Positional permutation while preserving internal sequence is therefore a biologically null manipulation, under which pathogenicity predictions should remain invariant.

Instead, sensitivity collapsed from 66.7% to 19.4%, demonstrating that Evo2’s variant scores are heavily driven by surrounding genomic context rather than intrinsic tRNA features. By contrast, structure-aware tools like RNAfold[33] evaluate the isolated transcript and remain inherently immune to such flanking artifacts.

### 5.5 NUMTs and the Limits of Contextual Reasoning

NUMTs test whether nuclear context enables Evo2 to distinguish non-functional pseudosequence from authentic mtDNA. Evaluated on a 5.8-kb, high-identity chromosome 1 NUMT (hg38: 629,079–634,924; spanning seven protein-coding genes, ND1-CO3 and 11 tRNAs), accurate sequence continuation required a narrow 500– 1,000 bp prefix window. Within this window, Evo2 favored the mitochondrial allele at nearly every divergent site, misreading the NUMT as functional mtDNA. Performance degradation beyond 1,000 bp mirrors flank-length sensitivities observed in pathogenicity scoring (optimal at 512 bp), indicating that context integration is governed by local window statistics rather than biological hierarchy.

These findings highlight broader vulnerabilities across pseudogenic regions: lacking an intrinsic mechanism to separate functional loci from non-functional paralogs, Evo2 defaults to functional sequence statistics whenever sequence similarity is high, confounding variant effect scores in pseudogene contexts.

### 5.6 Conservation, Evolutionary Constraint, and Gene Completion

Evolutionary constraint serves as a direct proxy for functional importance, leading to the expectation that high-performing genomic models assign greater likelihood to conserved positions. While Evo2 log-probabilities correlate strongly with PhyloP scores across *MT-RNR1* (Spearman *ρ* = 0.77), this relationship degrades locally: individual conservation peaks do not reliably match Evo2 likelihood maxima.

Similarly, highly mutation-intolerant loci should facilitate accurate sequence completion, yet Complex III (*CYTB*)—despite extreme disruption intolerance[41]—exhibited the lowest completion accuracy (85.0%) across all OXPHOS complexes. These discrepancies indicate that evolutionary signals are insufficiently resolved for fine-scale variant interpretation, aligning with recent evidence that Evo2 captures local sequence motifs while failing to preserve large-scale genomic correlations[42].

### 5.7 Comparison with custom-built tools

We benchmarked Evo2 against three purpose-built mtDNA predictors and one protein language model: MitoTIP[43], which scores tRNA variants from structural features; HmtVar[44], which integrates database and population evidence; APOGEE2[45], an ensemble trained on curated mtDNA missense data; and ESM-1v[46], a zero-shot protein language model, restricted to the variants scorable by every predictor compared (Table S5).

The custom tools win (Table 11), but APOGEE2 and MitoTIP were trained or calibrated on variant sets that likely overlap our labels, so their near-perfect scores are partly circular. The informative comparison is between the two genuinely zero-shot models, where the protein-level model substantially outperforms the DNA model on the same missense variants (AUROC 0.972 vs. 0.866), which is evidence that the limitation is not zero-shot prediction itself but what a nucleotide-level likelihood captures.

**Table 11.** Comparison of Evo2 with custom-built tools for variant-effect prediction in mtDNA. The variant numbers are explained in Table S5.

| **Variants** | **Predictor** | **AUROC (95% CI)** | **Balanced accuracy (95% CI)** |
|---|---|---|---|
| tRNA (n=81) | Evo2 (zero-shot) | 0.868 (0.778-0.940) | 0.823 (0.666-0.905) |
| tRNA (n=81) | MitoTIP | 0.988 (0.967-0.999) | 0.940 (0.844-0.974) |
| Missense (n=173) | Evo2 (zero-shot) | 0.866 (0.813-0.916) | 0.798 (0.712-0.850) |
| Missense (n=173) | ESM-1v (zero-shot) | 0.972 (0.947-0.988) | 0.923 (0.843-0.953) |
| Missense (n=173) | APOGEE2 | 1.000 (0.998-1.000) | 0.976 (0.882-0.990) |

This circularity may run deeper than training-set overlap. Computational predictions now carry enough weight in clinical classification to leave a detectable signature in ClinVar, concentrated in likely-benign calls[47], so variant labels cannot be treated as fully independent ground truth; a caveat that applies to our own benign set, drawn from ClinVar and ClinGen.

### 5.8 Plausible Reasons for Failure: Toward Biologically Grounded Foundation Models

Evo2’s autoregressive objective rewards local nucleotide prediction rather than mechanistic causality, conflating biological constraint with sequence correlation. As demonstrated by the tRNA permutation, NUMT, codon bias, and start/stop benchmarks, likelihood scoring inappropriately weights flanking context over intrinsic functional mechanisms. Because these failures persist despite a multi-scale architecture spanning codons to chromosomes[10], the primary limitation lies in the training signal rather than model capacity.

Variant interpretation is inherently hierarchical—resolving genomic compartment, coding consequence, and macromolecular impact in sequence. Lacking this hierarchy, autoregressive likelihoods cannot determine which context should govern scoring. Grounding foundation models will require multi-task objectives, functional fine-tuning, or hybrid architectures incorporating structure and conservation rather than simple unsupervised scaling. While Evo2 captures broad statistical regularities like transition bias and rank-level conservation, it lacks the causal constraint structure required for standalone clinical variant classification.

## 6 Conclusion & Future Work

Evo2 achieves competitive aggregate performance on mitochondrial variant-effect prediction (AUROC 0.927) yet systematically fails controlled biological tests: it mispredicts gene-specific codon usage, misclassifies canonical mitochondrial start codons as pathogenic, weights tRNA flanking context over internal sequence, misreads NUMTs as functional mtDNA, and completes the most constrained complex (Complex III) least accurately. These failures reflect a unifying flaw: predictions reflect sequence familiarity rather than biological constraint, causing standard aggregate metrics on imbalanced datasets to substantially overstate clinical readiness.

Moving forward, natural extensions include evaluating other foundation models and nuclear loci, fine-tuning on curated mitochondrial data, and incorporating Evo2 likelihood shifts (Δ*L*) into domain-specific metapredictors like APOGEE2[45]. Crucially, computational circularity in ClinVar[47] increasingly degrades the independence of supervised evaluation. The label-free, mechanism-grounded benchmarks established here—spanning codon usage, translation signals, structural invariance, and compartmental disambiguation—provide an essential blueprint for validating whether genomic models truly capture functional biology beyond unsupervised sequence scaling.

## 7 Reproducibility Statement

The mitochondrial reference is the revised Cambridge Reference Sequence (NC 012920.1); nuclear coordinates use GRCh38.p13/hg38. Variants were partitioned into calibration and test sets using a stratified 70/30 split (`sklearn.model selection.train test split`, `test size=0.3`, `random state=42`), yielding 551 calibration variants and 236 test variants. Variant labels were compiled from MITOMAP (confirmed pathogenic set, accessed August 2025) and ClinVar/ClinGen (pathogenic/likely pathogenic and benign sets, accessed August 2025); PhyloP conservation tracks were retrieved from the UCSC Table Browser (100-vertebrate alignment, accessed October 2025). Analyses used Python 3.12 with NumPy 2.3, pandas 2.2, SciPy 1.15, scikit-learn 1.7, and Biopython 1.85.

## 8 LLM Use Disclosure

ChatGPT was used for code debugging, boilerplate implementation and data-preprocessing assistance. All ChatGPT-generated code was thoroughly verified. Claude (Anthropic) was used for help with editing the manuscript and identifying gaps and inconsistencies in our narratives. The improvements were implemented by the authors after extensive discussions. All analyses, results, and scientific claims were reviewed and verified by the authors.

## Glossary

ENC: Effective Number of Codons is a summary statistic for a gene, measuring deviation of codon usage from the equal use of synonymous codons. ENC ranges from 20 (extreme bias) to 61 (unbiased, uniform usage of codons).
GC_3_: G/C fraction at the third nucleotide of codons
mtDNA: Mitochondrial DNA.
tRNA: Transfer RNA.
rRNA: Ribosomal RNA.
SNP: Single nucleotide polymorphism.
SNV: Single nucleotide variant.
indel: Insertion or deletion variant.
CRS: Cambridge Reference Sequence (human mtDNA reference, Genbank sequence name:NC 012920.1).
VEP: Variant Effect Prediction.
Synonymous mutation: A nucleotide change that does not alter the encoded amino acid.
Non-synonymous mutation: A nucleotide change that alters the encoded amino acid, including nonsense and missense mutations.
Missense mutation: A non-synonymous mutation that changes the encoded amino acid.
Nonsense mutation: A non-synonymous mutation resulting in a stop codon.
hg38: Human genome assembly GRCh38 (UCSC: hg38), the latest version of the human genome.
Heteroplasmy: Presence of more than one mitochondrial DNA genotype within a cell or individual.
NUMTs: Nuclear mitochondrial DNA segments; mitochondrial DNA fragments integrated into the nuclear genome.
MCC: Matthews Correlation Coefficient; a metric used to evaluate the performance of binary classification models (e.g., predicting whether a DNA variant is harmful or benign).
ROC: A Receiver Operating Characteristic (ROC) curve is a graphical plot that shows the performance of a binary classifier by plotting the true positive rate (sensitivity, y-axis) against the false positive rate (1 specificity, x-axis) across different classification thresholds. It illustrates the trade-off between detecting positives correctly and incorrectly labeling negatives as positives.
AUROC: Area Under the ROC curve; ranges from 0 to 1, is the numerical summary of a classifier’s performance across all possible classification thresholds. It represents the probability that the model ranks a randomly chosen positive example higher than a randomly chosen negative example. AUROC = 1.0 is perfect discrimination, AUROC = 0.5 is no better than random guessing, and AUROC ¡ 0.5 is worse than random (systematically misclassifying)

## Data and Code Availability

All code used in this study will be made available on GitHub.

## Supplementary Material

### S1 MELAS Mutations m.3243A*>*G and m.3252A*>*G: Structural Analysis

Both MELAS mutations act post-transcriptionally through intramolecular mechanisms independent of flanking genomic sequence (Figure S1), providing the biological premise for our tRNA permutation test. In *MT-TL1*, m.3243A*>*G impairs aminoacylation and post-transcriptional modification to disrupt translation[48], while m.3252A*>*G destabilizes the native cloverleaf fold (Table S1). A nearly threefold drop in minimum free energy (MFE) structure frequency and elevated ensemble diversity favor misfolded states, impeding Leucine charging and taurinomethyluridine modification, ultimately causing ribosomal stalling and proteotoxic OXPHOS stress above pathogenic heteroplasmy thresholds. Because neither mechanism involves surrounding genomic context, sensitivity to flanking sequence permutations indicates that a model captures local sequence regularities rather than true functional biology.

**Figure S1.**
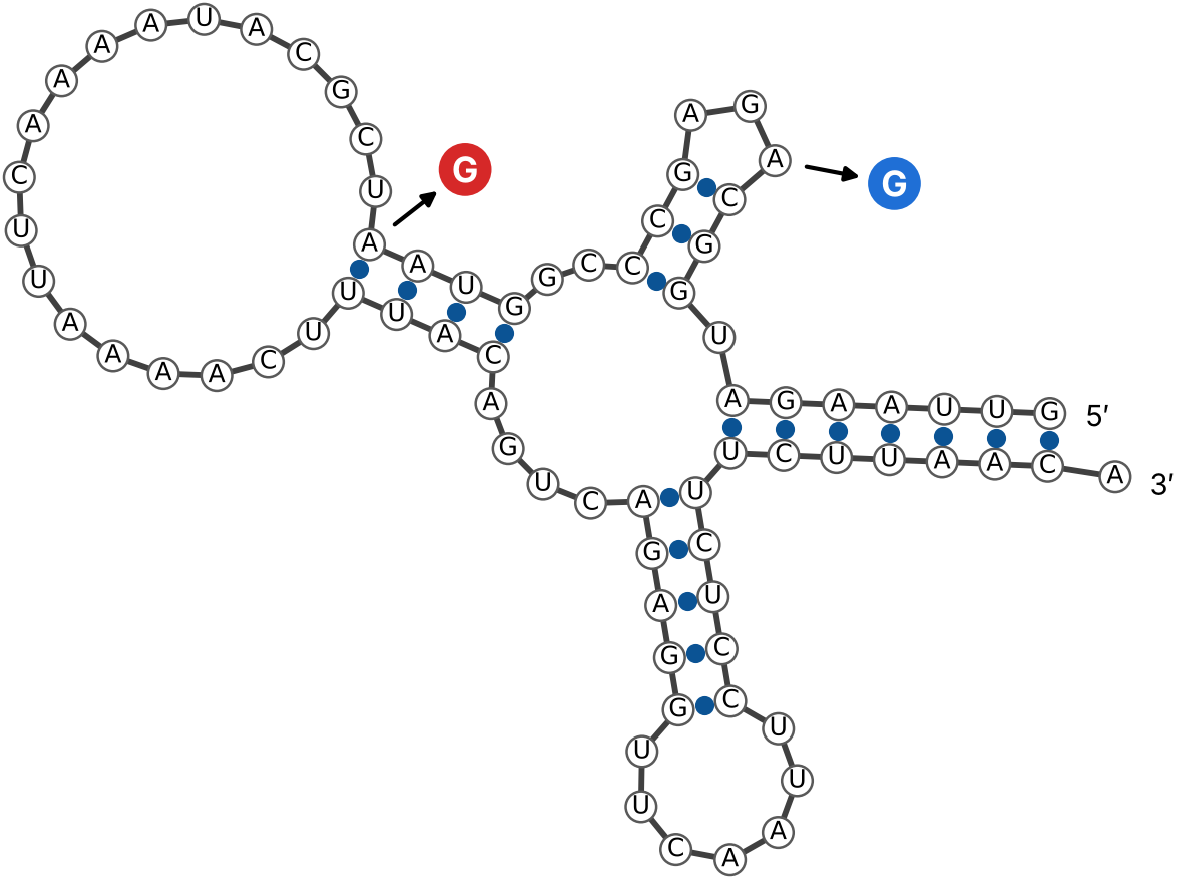
Secondary structure of the MT-TL1 leucine tRNA with MELAS mutation sites highlighted. The m.3243A*>*G site is shown in blue (the more common MELAS cause) and m.3252A*>*G in red (rarer). Secondary structures were generated with RNAfold[33]. Although the predicted folds appear similar, the two mutations act through distinct mechanisms: m.3243A*>*G impairs aminoacylation, while m.3252A*>*G destabilizes the binding to the U (3272) that is 20 nucleotides away, thereby destabilizing the cloverleaf structure leading to functional impairment (native fold Table S1).

**Table S1.**
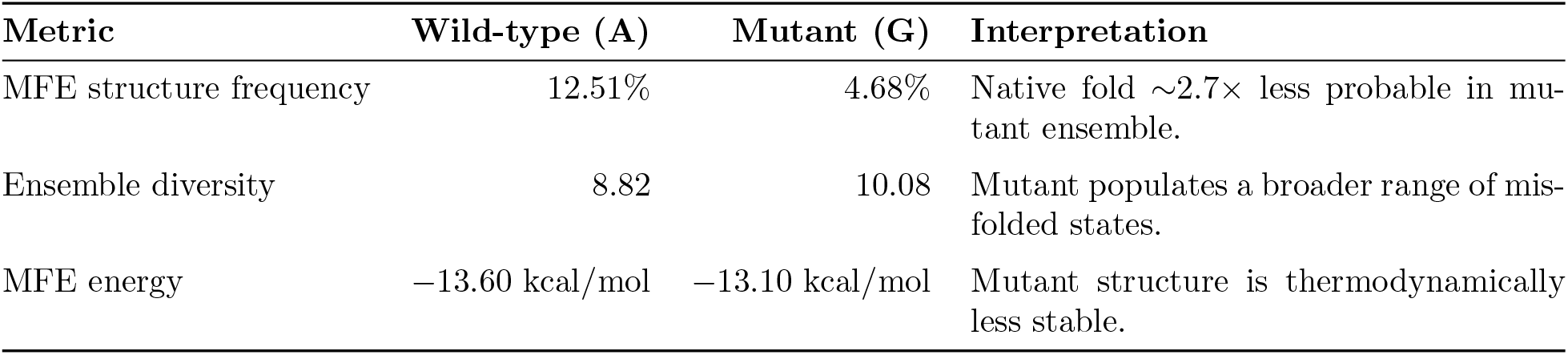
Thermodynamic comparison of wild-type and MELAS mutant (m.3252A*>*G) MT-TL1 RNA folding (Figure S1).

| **Metric** | **Wild-type (A)** | **Mutant (G)** | **Interpretation** |
|---|---|---|---|
| MFE structure frequency | 12.51% | 4.68% | Native fold ∼2.7× less probable in mu-tant ensemble. |
| Ensemble diversity | 8.82 | 10.08 | Mutant populates a broader range of mis-folded states. |
| MFE energy | −13.60 kcal/mol | −13.10 kcal/mol | Mutant structure is thermodynamically less stable. |

### S2 Confusion Matrix and Its Relation to ROC/AUROC

For binary variant-effect prediction (VEP), variants are classified as pathogenic (deleterious) or benign (neutral). Each prediction falls into one of four categories, shown in this *Confusion Matrix* :

**Table S2.** Confusion matrix for binary variant-effect prediction. Quantities calculated from this are shown in Table S3

|  | **Predicted Pathogenic** | **Predicted Benign** |
|---|---|---|
| **True Pathogenic** | True Positive (**TP**) pathogenic cor-rectly predicted as pathogenic. | False Negative (**FN**) pathogenic wrongly predicted as benign |
| **True Benign** | False Positive (**FP**) benign wrongly pre-dicted as pathogenic | True Negative (**TN**) benign correctly pre-dicted as benign |

From this matrix standard performance metrics can be calculated as shown in Table S3.

In variant interpretation, error types carry distinct biological consequences: false negatives understate pathogenic risk, whereas false positives artificially inflate disease burden. Because continuous variant-effect scores require thresholding to generate discrete confusion matrices (TP, FP, TN, FN), single summary statistics can obscure clinical trade-offs.

**Table S3.** Performance metrics derived from the confusion matrix (Table S2)

| **Metric** | **Formula** |
|---|---|
| Accuracy | $\frac{TP+TN}{TP+TN+FP+FN}$ |
| Sensitivity (Recall) | $\frac{TP}{TP+FN}$ |
| Specificity | $\frac{TN}{TN+FP}$ |
| Precision (Positive Predictive Value) | $\frac{TP}{TP+FP}$ |

#### Connection to ROC and AUROC, and Imbalance Limitations

The Receiver Operating Characteristic (ROC) curve evaluates sensitivity versus (1-specificity) across all cutoffs, with the Area Under the Curve (AUROC) measuring global ranking ability. In variant-effect prediction, however, severe class imbalance (benign ≫ pathogenic) allows AUROC to remain high even when performance on the pathogenic minority class is inadequate. A strong AUROC often masks unacceptable false-negative or false-positive rates at specific clinical operating points; thus, it is essential to have threshold-specific confusion matrices or precision-recall curves which focus explicitly on the pathogenic class.

#### Vulnerabilities of MCC Under Severe Imbalance

The Matthews Correlation Coefficient (MCC)[49],

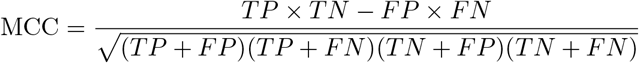

represents the Pearson correlation for binary variables. Under extreme skew, the minority-class denominator terms shrink, blunting the metric’s sensitivity to pathogenic classification errors—the exact domain of clinical interest.

### S3 Forward vs. backward completion

Per-species breakdown of the directional comparison reported in Section 4.5.

**Figure S2.**
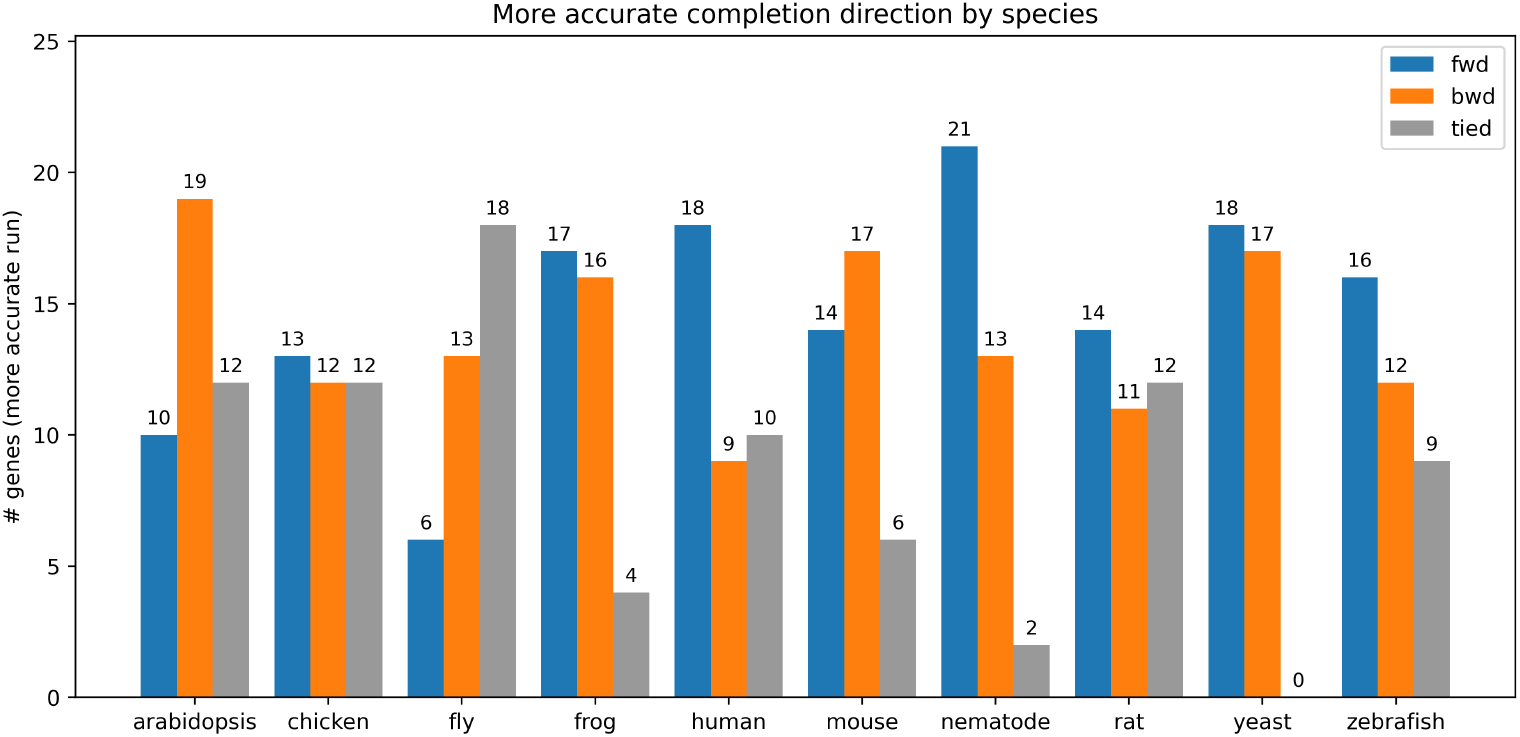
Forward vs. backward completion accuracy by species. For each gene, the run (forward or backward) that completed the masked middle third more accurately was recorded; tied genes had equal accuracy in both runs. Across 371 genes, the forward run was more accurate for 147, the backward run for 139, and the two tied for 85.

### S4 Start and Stop Codons for 13 Protein-Coding Mitochondrial Genes

**Table S4.** Start and stop codons for the 13 human mitochondrial protein-coding genes. ^*†*^Incomplete stop codons. The nuclear start codons are usually AUG along with rarer variants CUG, GUG, or UUG. Nuclear stop codons are UAA, UAG, and UGA. Human mtDNA start/stop codons are more variable; start codons are AUG (9), AUA (3), and AUU (1), allowing more variability while stop codons are UAA(3), UAG(2), AGA(1), AGG (1). The remaining 6 genes end in an incomplete stop codon, which is completed to UAA post-transcriptionally by polyadenylation of the mRNA. This diversity of non-canonical start and stop codons reflects the evolutionary divergence of the vertebrate mitochondrial and nuclear genetic codes.

| Type | ND1 | ND2 | COX1 | COX2 | ATP8 | ATP6 | COX3 | ND3 | ND4L | ND4 | ND5 | ND6 | CYTB |
|---|---|---|---|---|---|---|---|---|---|---|---|---|---|
| Start codon | AUA | AUU | AUG | AUG | AUG | AUG | AUG | AUA | AUG | AUG | AUA | AUG | AUG |
| Stop codon | UA<sup>†</sup> | U<sup>†</sup> | AGA | UAG | UAG | UAA | U<sup>†</sup> | U<sup>†</sup> | UAA | U<sup>†</sup> | UAA | AGG | U<sup>†</sup> |

### S5 Datasets Used in Each Analysis

**Table S5.**
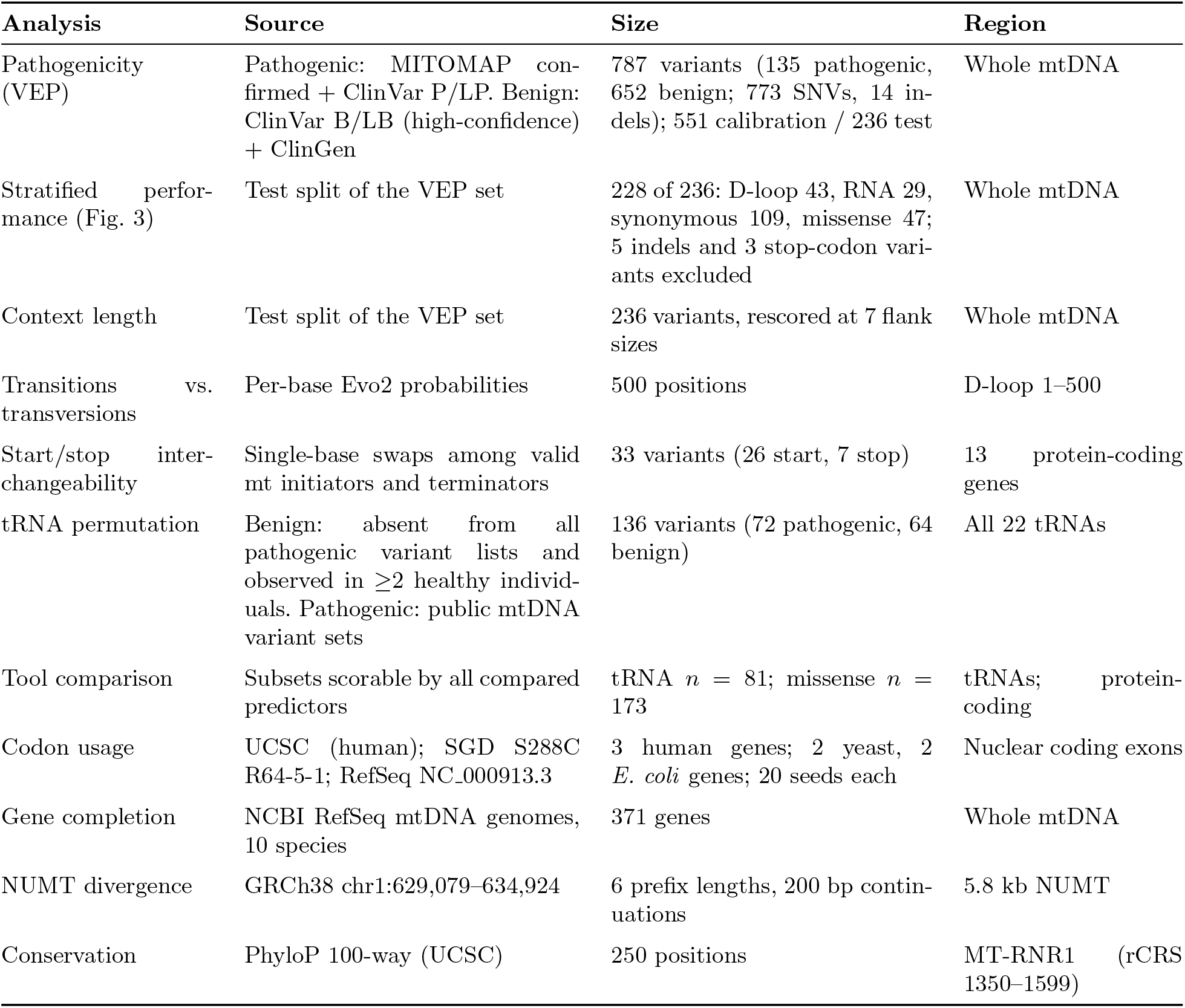
Datasets used in each analysis. Variant counts, sources and genomic scope for every test reported in this study.

| **Analysis** | **Source** | **Size** | **Region** |
|---|---|---|---|
| Pathogenicity (VEP) | Pathogenic: MITOMAP con-firmed + ClinVar P/LP. Benign: ClinVar B/LB (high-confidence) + ClinGen | 787 variants (135 pathogenic, 652 benign; 773 SNVs, 14 in-dels); 551 calibration / 236 test | Whole mtDNA |
| Stratified perfor-mance (Fig. 3) | Test split of the VEP set | 228 of 236: D-loop 43, RNA 29, synonymous 109, missense 47; 5 indels and 3 stop-codon vari-ants excluded | Whole mtDNA |
| Context length | Test split of the VEP set | 236 variants, rescored at 7 flank sizes | Whole mtDNA |
| Transitions vs. transversions | Per-base Evo2 probabilities | 500 positions | D-loop 1–500 |
| Start/stop inter-changeability | Single-base swaps among valid mt initiators and terminators | 33 variants (26 start, 7 stop) | 13 protein-coding genes |
| tRNA permutation | Benign: absent from all pathogenic variant lists and observed in ≥2 healthy individ-uals. Pathogenic: public mtDNA variant sets | 136 variants (72 pathogenic, 64 benign) | All 22 tRNAs |
| Tool comparison | Subsets scorable by all compared predictors | tRNA *n* = 81; missense *n* = 173 | tRNAs; protein-coding |
| Codon usage | UCSC (human); SGD S288C R64-5-1; RefSeq NC_000913.3 | 3 human genes; 2 yeast, 2 *E. coli* genes; 20 seeds each | Nuclear coding exons |
| Gene completion | NCBI RefSeq mtDNA genomes, 10 species | 371 genes | Whole mtDNA |
| NUMT divergence | GRCh38 chr1:629,079–634,924 | 6 prefix lengths, 200 bp contin-uations | 5.8 kb NUMT |
| Conservation | PhyloP 100-way (UCSC) | 250 positions | MT-RNR1 (rCRS 1350–1599) |

## Notes

### Competing Interest Statement

The authors have declared no competing interest.

### Summary of Updates

This revision adds new analyses and tightens existing ones. We now compare Evo2 directly against purpose-built mtDNA predictors (MitoTIP, HmtVar, APOGEE2) and a zero-shot protein language model (ESM-1v), and find that the protein-level model substantially outperforms Evo2 on the same missense variants. The transition/transversion analysis has been replaced with a per-base measurement of Evo2's predicted nucleotide probabilities across the D-loop, which assays the model's mutational prior directly rather than through an ascertainment-biased variant set. The codon-usage benchmark now extends to highly expressed genes in S. cerevisiae and E. coli, testing translational selection alongside compositional bias, and we add a context-length sensitivity analysis with thresholds recalibrated at each flank size. Variant categories in the stratified analysis were refined and the associated figure and statistics regenerated; the forward/backward completion comparison now uses accuracy rather than log-likelihood. We state two limitations explicitly: conservation at start/stop positions is a partial confound, and overlapping reading frames mean some start/stop variants may be correctly called pathogenic. A new supplementary table specifies the source, size and scope of every dataset used, and we cite recent work showing that variant-effect predictors are already shaping the ClinVar classifications used to train and evaluate them, which is a circularity that motivates the label-free benchmarks developed here. Figures and text were revised throughout for clarity.

